# A ribosome exit tunnel component is required for robust expression of the *Drosophila* hox gene, *abdominal A*

**DOI:** 10.64898/2026.10.23.761343

**Authors:** Matthew R. McFarland, Ratna Tripathy, Jennifer K. Bergman, Michael Wood, Andrew D. Renault

## Abstract

Ribosomes, the molecular machines responsible for protein synthesis, are fundamental to all cellular life. Although the higher-level architecture of ribosomes is conserved, a small subunit acting as the mRNA decoding site and a large subunit that catalyses peptide bond formation, there are differences in the numbers of components of each of the subunits between the 3 domains of life. Despite their well-established role in translating genetic information into functional proteins, many complexities in ribosome biology and functionality are only just being revealed. Defects in ribosomal components contribute to a class of disorders termed ribosomopathies that come with a range of symptoms and severities. Here we show that a mutation in the *Drosophila* ribosomal protein RpL39 causes an anterior-posterior embryonic patterning defect through impaired accumulation of the hox protein Abdominal-A (abdA). Deletions uncovering subunits of the Nascent peptide-Associated Complex (NAC) and Ribosome-Associated Complex (RAC) act as genetic enhancers of the RpL39 mutation leading us to speculate that RpL39 forms a relay between the nascent peptide in the exit tunnel and the exterior surface of the ribosome, modulating the interaction with these ribosomally-associated complexes. This work enhances our knowledge of the roles played by individual ribosome components and feeds into our understanding of the mechanisms of ribosome quality control.

## Introduction

Ribosomes are fundamental and conserved cellular machines that translate mRNAs into amino acid sequences. In all species ribosomes are composed of a small subunit which acts as the mRNA decoding site, and a large subunit which catalyses peptide bond formation. As it is assembled, the nascent peptide passes through the exit tunnel within the large subunit that links the ribosome’s factory floor to the cellular environment (Kramer et al., 2009). Folding of the nascent chain occurs co-translationally, with smaller protein domains able to fold within the confines of the tunnel itself (Nilsson et al., 2015). Ribosomally-associated chaperones, such as the Ribosome- and Nascent peptide-Associated Complexes (RAC and NAC respectively) of eukaryotes, interact near the tunnel exit and assist with folding and localisation of the emerging peptide (Deeng et al., 2016; Koplin et al., 2010; Pech et al., 2010; Peisker et al., 2008).

Despite the highly conserved nature of many of the ribosome’s features including its fundamental function, structure and sequence of constituent proteins, there are several ways through which the ribosome itself can contribute to selective mRNA translation in different situations. One of these is via heterogeneity in ribosome composition. In many species, heterogeneity results from the presence of additional ribosomal protein paralogues (Gerst, 2018). The incorporation of an alternative ribosomal protein generally occurs in a particular tissue or in response to particular intra- or extra-cellular conditions and for some cases this has been shown to alter the spectrum of translated mRNAs (Shi et al., 2017; H. W. Yang et al., 2019). Another mechanism involves the presence or absence of ribosomal components, often in response to cellular stress (Y.-M. Yang & Karbstein, 2026).

Selective translation can also be achieved through a function that can be attributed to a core ribosomal protein. For example, haploinsufficiency of RpL38 in mouse leads to tissue-specific patterning defects during embryonic development. This is caused by a lack of translation of several *hox* genes (Kondrashov et al., 2011). These *hox* mRNAs contain internal ribosome entry sites (IRESs) in their 5’ UTRs that require RpL38 for their utilisation (Xue et al., 2015). These mRNAs also contain inhibitory elements in their 5’ UTRs that block normal cap-dependent translation, meaning they are reliant on their IRESs for robust translation (Xue et al., 2015).

Mutations in many ribosomal proteins are causative for a variety of human diseases, collectively termed ribosomopathies. Given the essential role of ribosomes, surprisingly, these can have a range of symptoms and severities. The best characterised of these is Diamond-Blackfan Anaemia (DBA), which is caused by mutations in one of at least 15 ribosomal proteins. This results in a low red cell count (anaemia) accompanied in approximately half of cases by a range of congenital malformations, the type and severity of which appear to be dependent on the affected ribosomal protein (Gazda et al., 2008; Horos et al., 2012). Another ribosomopathy, Isolated Congenital Asplenia, results from haploinsufficiency of ribosomal protein SA and leads to a defect in splenic development but with no other observable abnormalities (Bolze et al., 2013). Whilst these phenotypes may result from reduced levels of fully functioning ribosomes or lack of specifc ribosomal functions, some phenotypes are secondary effects caused by activation of the p53 pathway by imbalanced ribosomal protein levels leading to cell cycle arrest and apoptosis (Bai et al., 2014; Jones et al., 2008; Xiong et al., 2014).

The surprising diversity of known ribosomal protein mutant phenotypes and their involvement in disease make a strong argument for building a comprehensive picture of the roles for ribosomal proteins, most usefully in a model organism. In *Drosophila*, most null mutations in ribosomal proteins are haploinsufficient, meaning that they are not viable in the heterozygous condition. Weaker loss of function mutations, however, are heterozygous viable but show dominant ‘Minute’ phenotypes, characterized by short adult macrochaetae (sensory bristles) and delayed development (Bridges & Morgan, 1923; Marygold et al., 2007). This is believed to be caused by reduced overall rates of protein synthesis that particularly affect cells which must produce a large amount of protein in a short developmental time window (Lambertsson, 1998). Interestingly these mutations can also lead to cell competition (Morata & Ripoll, 1975). In genetically mosaic animals, clones of cells heterozygous for ribosomal protein mutations die when in contact with wild type cells. This allows the developing organism to eliminate those cells with an incomplete functional complement of ribosomal protein genes.

In this study, we identify a mutation in the *Drosophila* ribosomal protein *RpL39* (also known as eL39). This mutation is not dominant but, surprisingly, is a maternal effect and causes an anterior-posterior embryonic patterning defect. This mutation specifically impairs the translation of the hox protein Abdominal-A (abdA) due to a sequence feature within the AbdA N-terminal domain. We further show that subunits of the Nascent peptide-Associated Complex (NAC) and Ribosome-Associated Complex (RAC) act as genetic enhancers of the RpL39 mutation. From these observations, and the protein’s position in the vestibule of the ribosome exit tunnel (Nissen et al., 2000; Zhang et al., 2013), we speculate that RpL39 forms a relay between the nascent peptide in the exit tunnel and the exterior surface of the ribosome, modulating the interaction with these ribosomally-associated complexes.

## Results

### RpL39^A44^ causes defective gonad formation in Drosophila

In a forward genetic screen designed to identify genes important for development of the embryonic gonad, we identified a mutant herein termed *A44*. This mutation shows a maternal effect, with mothers homozygous for *A44* producing eggs that show defects in germ cell migration (Fig. 1A-C, G-I) regardless of whether the paternal flies are wild-type or *A44* mutants themselves (data not shown).

**Figure 1:**
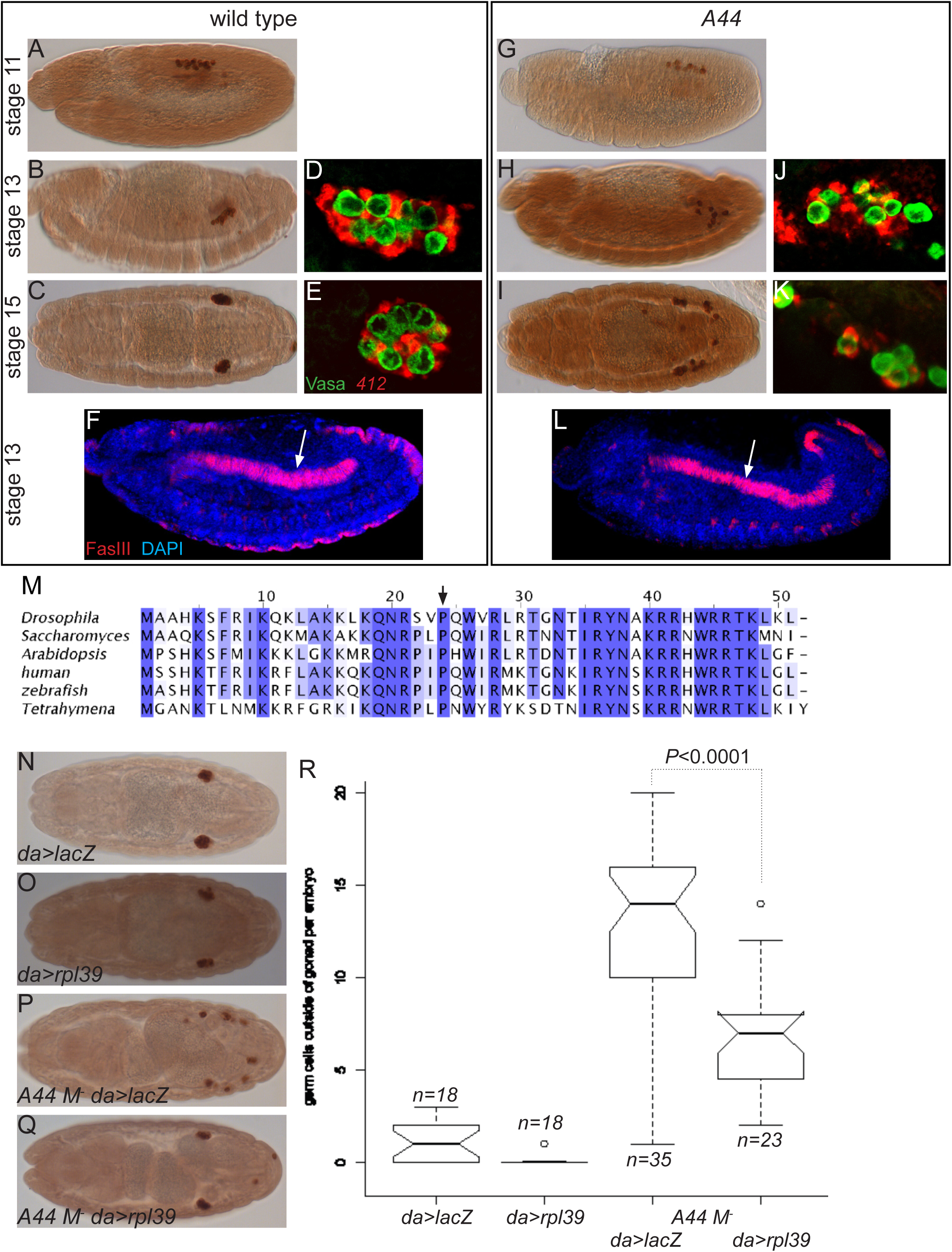
A44 is a mutant in a conserved residue of Drosophila RpL39 which causes gonad formation defects. (A-C; G-H) Lateral (A,B,G,H) and dorsal views (C,I) of embryos from wild type (A-C) and *A44* M^-^ mutants (G-I) stained with an antibody against Vasa to highlight the germ cells showing the mis-migration occurring from stage 13 onwards in *A44* mutants. (D,E,J,K) Embryos stained with antibodies against Vasa (green) and hybridised with the *412* riboprobe which labels the SGPs (red) showing that the germ cell mis-migration in *A44* mutants (J,K) results from a failure of the SGPs to form a single cluster and then compact as in wild type (D,E). (F, L) Lateral views of a stage 13 wild type (F) and *A44* mutant (L) embryo stained with an antibody against FasIII to highlight the trunk visceral mesoderm (arrows) which is continuous in both wild type and *A44* mutants, indicating that other mesodermally derived tissues are not affected in *A44* mutants. (M) Multiple sequence alignment of *Drosophila* RpL39 with homologues from *Saccharomyces cerevisiae* (Accession: NP_012346.1), *Arabidopsis thaliana* (NP_567886.1), Humans (NP_000991.1), zebrafish (NP_001002078.1) and Tetrahymena (XP_001028051.2). (N-Q) Dorsal views of stage 16 embryos stained with an antibody against Vasa to highlight the germ cells. Ubiquitous over-expression (using *daughterless* Gal4) of LacZ as a control (N) or RpL39 (O) does not disrupt gonad formation. Embryos from *A44* homozygous mothers over-expressing RpL39 (Q) show reduced numbers of mis-migrated germ cells compared to those over-expressing LacZ (P). (R) Box plot showing the number of germ cells outside the gonad per embryo for the genotypes depicted in N-Q. *P* value from Student’s t test.

As the germ cell migration in *A44* remained similar to that of wild-type until stage 13 onwards, we hypothesised this defect may be the result of changes in the number or behaviour of the somatic gonadal precursor cells (SGPs), which begin to ensheathe the germ cells at this stage (Boyle & DiNardo, 1995). To investigate this possibility, we labelled SGPs in *A44* mutant embryos using a riboprobe to the *412* retrotransposon, a marker of SGP cell identity (Brookman et al., 1992) (Fig. 1J, K). When compared to wild-type embryos (Tripathy et al., 2014) (Fig. 1D, E), we found that the SGPs failed to coalesce correctly around the germ cells. This defect is specific to SGPs with the localisation of other mesodermally-derived tissue, such as the visceral mesoderm (Fig. 1F, L), appearing to be unaffected in *A44* embryos.

Together, these results suggest that while *A44* is a maternal effect mutation and the affected gene likely acts at the very early stages of development, the resulting defect only becomes apparent later in embryogenesis.

The causative mutation in *A44* was identified through a combination of recombination mapping and whole genome sequencing (Fig. S1 and Supplementary Material) as a SNP in the *RpL39* gene that results in the substitution of a conserved proline residue with serine (P24S) (Fig. 1M). *RpL39* encodes for a subunit of the 60S ribosome, where it forms part of the vestibule of the ribosome exit tunnel (Zhang et al., 2013). To determine if this SNP is responsible for the observed defects in gonad formation, we attempted to rescue the phenotype by expressing wild-type *RpL39* using the *GAL4*-UAS system with the early ubiquitous driver *daughterless>GAL4* (*daGAL4*). Indeed, expression of *RpL39*, but not a control protein (*lacZ*), significantly reduced the number of germ cells outside the gonad in *A44* embryos (Fig. 1N-R). We therefore conclude that the defects seen in the *A44* mutants are the result of a point mutation in the *RpL39* gene (hereafter *RpL39^A44^*).

### RpL39^A44^ phenocopies mutants in AbdA

Mutants in over fifty zygotic genes are known to affect the specification or behaviour of the SGPs (Jemc, 2011). To narrow down which of these genes might be being affected by *RpL39^A44^,* we examined *RpL39^A44^*for more general patterning defects using the denticle belts of the larval cuticle as a marker. The anterior abdominal denticle bands of *RpL39^A44^* larvae were noticeably narrowed compared to those of their wild-type counterparts (Fig. 2A, B). This pattern phenocopies mutants of the hox gene *abdominal A* (*abdA*) (Fig. 2C), and is distinct from other hox gene mutants, such as *abdominal B* (*abdB*) (Fig. 2D). The expression domain of *abdA* (parasegments 7-12) includes the region in which the embryonic gonad is specified (parasegments 10-12) (Brookman et al., 1992; Karch et al., 1990), and *abdA* mutations display defects in germ cell migrations and SGP specification and behaviour (Moore et al., 1998). Together, these data suggest that *RpL39^A44^* affects gonad development through impacting *abdA* activity.

**Figure 2:**
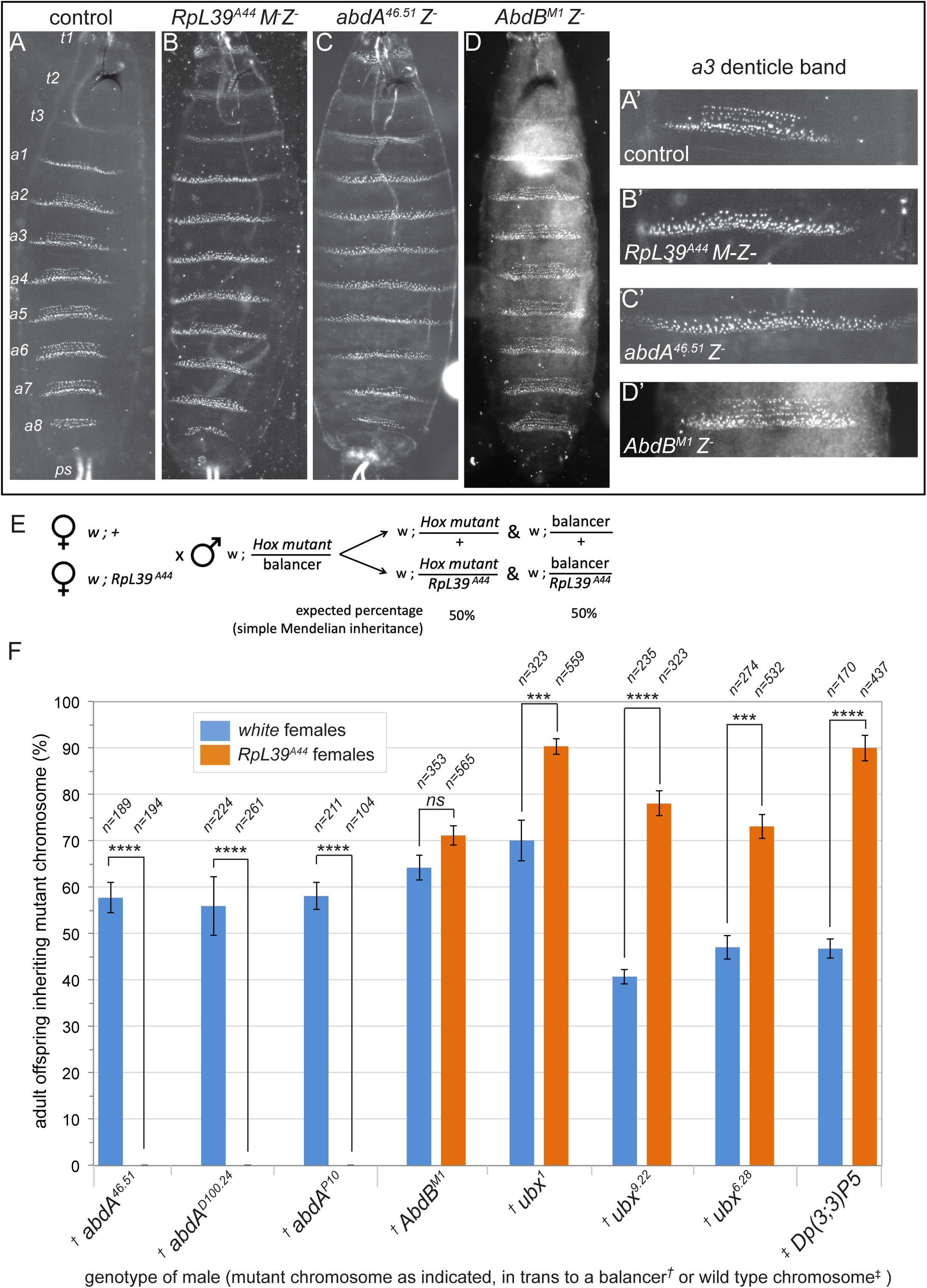
RpL39^A44^ phenocopies and genetically interacts with abdA. (A-D) Cuticles of control *white* (A, A’), *RpL39^A44^* (B, B’), *abdA^46.51^* (C,C’) and *AbdB^M1^* (D,D’) larvae showing how the denticle band defects of *RpL39^A44^* phenocopies those of *abdA* but not *AbdB* mutants. The denticle bands of abdominal segments 2 – 7 (a2 – a7) are thinner and elongated in both *A44* and *abdA* mutants which is especially apparent in the enlargements of a3 (A’-D’). In contrast, *AbdB* mutants lack posterior spiracles (ps), and the denticle bands of abdominal segments 5 – 8 (a5 – a8) are a3-like, appearing highly trapezoidal. t1 – 3 indicate the thoracic denticle bands. (E) Crossing scheme to test for genetic interactions between *RpL39^A44^* and Hox genes. *white* control females (upper line) and *RpL39^A44^* females (lower line) were mated to males carrying a Hox gene mutation in trans to a dominantly marked balancer chromosome (in this example located on chromosome II). Numbers of offspring inheriting the Hox gene mutant chromosome versus the balancer were scored. (F) Graph showing the percentage of offspring inheriting the Hox gene mutant chromosome from the crosses shown in E. Significance shown based on Student’s t test (ns P > 0.05; * P ≤ 0.05; **P ≤ 0.01; *** P ≤ 0.001; **** P ≤ 0.0001).

To test this hypothesis, we tested for a genetic interaction between *RpL39^A44^* and *abdA*. We mated homozygous *RpL39^A44^* females (or *white* females as a control) to males carrying independent *abdA* mutations in *trans* to a dominantly marked balancer chromosome and scored the adult offspring that emerged for inheritance of the mutant chromosome versus the balancer (Fig. 2E). Strikingly, none of the emerged adult offspring of *RpL39^A44^* females inherited the mutant chromosome for any of the three *abdA* alleles tested, compared with approximately 50% inheritance for *white* control females, as expected (Fig. 2F). Therefore, possessing only one genomic copy of *abdA*, which is sufficient for survival of embryos from wild type females, is insufficient in embryos from *RpL39^A44^* mutant mothers.

We examined whether *RpL39^A44^* also showed an interaction with *abdA* in SGP behaviour. We found that a reduction in *abdA* genomic copy number enhanced the gonad formation defects in embryos from *RpL39^A44^* mutant mothers (Fig. S2). Together, these data suggest that *abdA* mutants are strong genetic enhancers of *RpL39^A44^*-induced lethality and gonad formation defects.

We next asked if the reciprocal interaction also occurs, whether additional copies of *abdA* can suppress the effects of *RpL39^A44^*. We crossed homozygous *RpL39^A44^*females to males carrying a dominantly marked chromosome in which the entire bithorax hox gene complex (BX-C), including *abdA*, is duplicated (*Dp(3;3)P5 Sb^1^*) in *trans* to a wild-type chromosome. The percentage of resulting adult offspring inheriting the duplication was significantly increased in the case of *RpL39^A44^* females versus *white* control females (Fig. 2F). Possession of an additional genomic copy of *abdA* therefore increases the survival of offspring from *RpL39^A44^* females.

To examine the specificity of the genetic interaction between *RpL39^A44^* and *abdA*, we crossed *RpL39^A44^* females to males carrying alleles of the other bithorax complex genes, *Ultrabithorax* (*Ubx*) and *abdB*, which are expressed immediately anterior to, and posterior to *abdA* respectively. In the case of *abdB*, the percentage inheritance of the *abdB^M1^* mutant chromosome was not significantly different between control and *RpL39^A44^* females (Fig. 2F). However, when crossed to *Ubx* mutant males, we saw a significant increase in inheritance of the mutant chromosome in the offspring of *RpL39^A44^* females versus controls with all three alleles tested (Fig. 2F). From this, we conclude that *Ubx* is a genetic suppressor of *RpL39^A44^*-induced lethality.

### RpL39^A44^ reduces AbdA protein levels

As RpL39 forms part of the ribosomal exit tunnel, one possible explanation for the phenocopy and genetic interaction with *abdA* mutants is that the *RpL39^A44^* mutation impairs *abdA* translation such that it sits at the threshold for embryonic development. Increases and decreases in *abdA* genomic copy number would cause the numbers of embryos reaching this threshold to increase and decrease respectively.

To test this hypothesis, we used whole mount immunofluorescence to examine the AbdA protein expression pattern and levels in wild type and *RpL39^A44^* embryos. AbdA expression is first detectable at stage 8-9 and by stage 14 AbdA is expressed in the epidermis of parasegments 7 to 12 (Gummalla et al., 2012; Karch et al., 1990). Although the AbdA expression pattern in *RpL39^A44^*was identical to wild type, and the protein was predominantly nuclear, fluorescence intensity levels were markedly reduced in *RpL39^A44^* embryos compared to controls (Fig. 3A, B).

**Figure 3:**
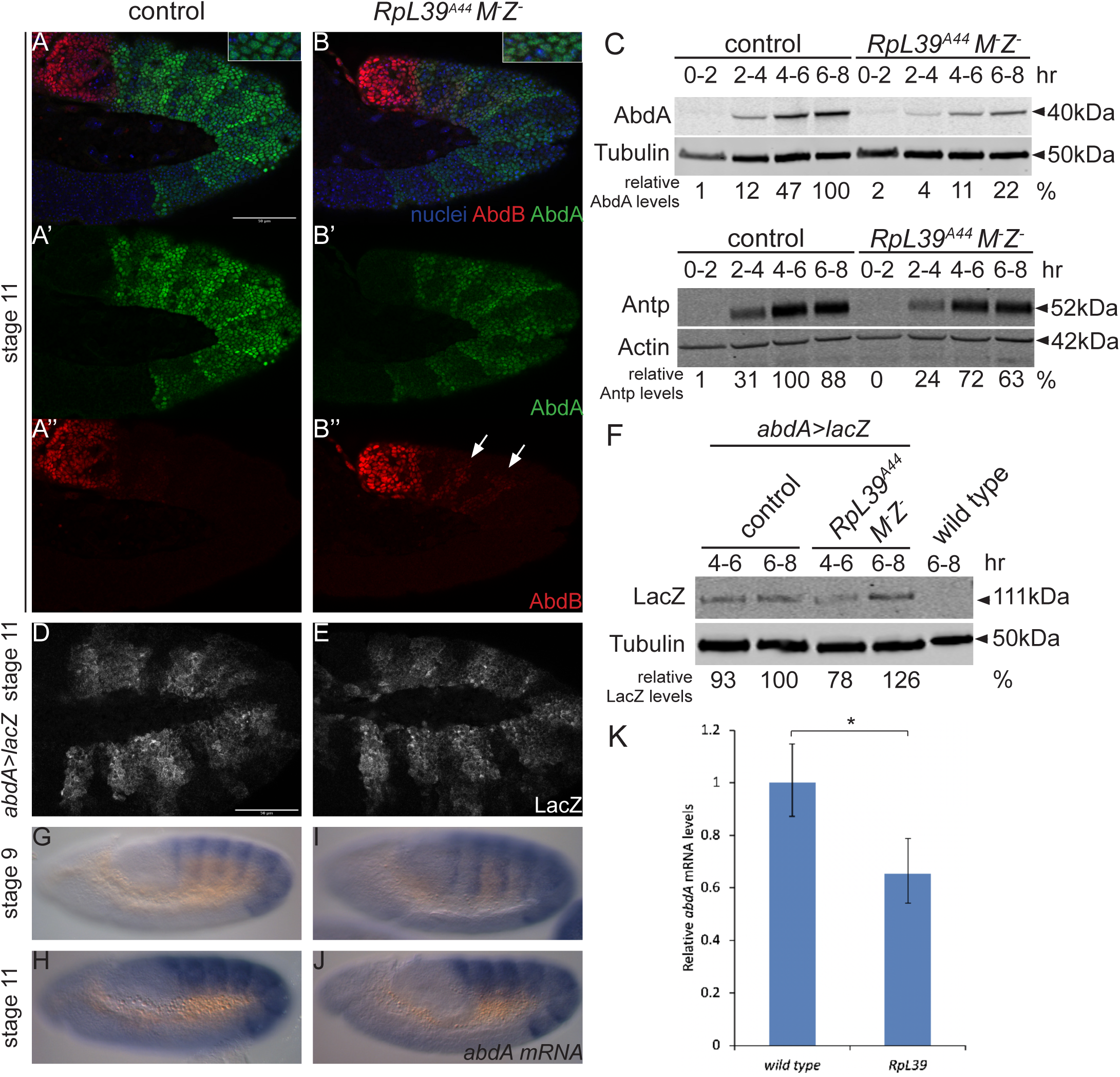
AbdA protein but not mRNA levels are reduced in RpL39^A44^ mutants. (A,B) Lateral views of stage 11 control (*w*) (A) and *RpL39^A44^* M^-^Z^-^ (B) embryos stained with antibodies against AbdA (green), AbdB (red) and DAPI to highlight nuclei (blue) showing reduced AbdA but not AbdB levels in the mutant. Insets show a region at increased magnification highlighting the overlap between AbdA and DAPI. (C) Western blots of aged embryos from control (*w*) and *RpL39^A44^* flies. In both cases AbdA levels increase as development proceeds, however levels are consistently lower in *RpL39^A44^* embryos (upper blot). Mean level of AbdA in 4-6hr *RpL39^A44^* M^-^Z^-^ embryos was 29% (n=4, s.d.=20%) compared to controls. To demonstrate that *RpL39^A44^* M^-^Z^-^ embryos were developing at the same rate as wild type, the same protein extracts were tested for the increase of expression of another Hox gene, Antp (lower blot). Relative levels indicate AbdA or Antp band intensity normalized using the loading control (tubulin or actin respectively) and then shown as a percentage relative to the AbdA or Antp band of highest intensity. (D-E) Lateral views of stage 11 embryos from wild type (D) or *RpL39^A44^* (H) females crossed to males containing an *abdA* driven *lacZ* construct stained with an antibody against LacZ showing identical patterns and staining intensities. (F) Western blots probed with the indicated antibodies of aged wild type embryos (*w*) and control and *RpL39^A44^* M^-^Z^-^ embryos which carry a construct that expresses *lacZ* from the *abdA* promotor (from a cross of *w* and *w; RpL39^A44^* mothers respectively mated to *RpL39^A44^; MKRS/TM6abdA>lacZ* males) showing that β-galactosidase levels are not reduced by the *RpL39^A44^*mutation. Mean levels of β-galactosidase in 6-8hr *RpL39^A44^*M^-^Z^-^ embryos was 135% (n=3, s.d.=29%) compared to controls. (G-J) Lateral views of stage 9 (G,I) and 11 (H,J) wild type (G,H), and *RpL39^A44^* (I,J) embryos hybridized with an *abdA* riboprobe showing identical patterns and staining intensities. (K) Graph showing relative *abdA* mRNA levels by quantitative RT-PCR in 6-8 hr wild type and *RpL39* embryos. Data is mean from 3 replicates, error bars show relative quantification (RQ) min and max values. The reduction in *abdA* expression in *RpL39* mutant is significant (Student’s t test, p=0.014).

To determine whether other Hox genes were also affected, we examined the expression pattern of AbdB. In contrast to AbdA, we found that AbdB levels were not reduced in *RpL39^A44^* embryos compared to wild type, indicating that the effects of *RpL39^A44^* on AbdA are specific (Fig. 3A,B). We did observe an expansion in the extent of AbdB expression in *RpL39^A44^* embryos, in the form of additional stripes of AbdB expression inside the expression domain of AbdA (arrows in Fig. 3B’’). The prevailing dogma (often referred as to as “posterior dominance/prevalence”) states that posterior Hox gene (in this case *AbdB*) normally represses the expression of the immediately adjacent anterior Hox gene (in this case *abdA*) (Karch et al., 1990). However, there is evidence that this functional dominance is somewhat reciprocal and is in part due to the Hox genes competing to positively or negatively regulate the same targets in an opposing manner (Capovilla & Botas, 1998). To test this hypothesis, we examined AbdB expression in *abdA^46.51^* embryos and observed expansion of AbdB expression compared to sibling control embryos, identical to that seen in *RpL39^A44^* embryos (Fig. S3). We also examined AbdA levels in *Ubx* mutants to test if the genetic suppression of *RpL39^A44^* induced lethality by *Ubx* might result from Ubx normally repressing AbdA expression in early embryos. We were unable to detect any difference in AbdA levels in Ubx1 mutants compared to sibling controls (Fig. S3).

To quantify the reduction in AbdA levels observed by immunofluorescence, we performed quantitative western blotting on aged embryos. In wild type embryos, AbdA levels rapidly increase from 2-4 hrs to 6-8 hrs (Fig. 3C, upper blot). In *RpL39^A44^* embryos, AbdA levels also increase, but the level at each time point is less than wild type (Fig. 3C, upper blot). As a control, we examined the levels of another Hox gene, Antennapedia (Antp), and found that the pattern of expression of Antp levels increased similarly over time in both wild-type and *RpL39^A44^*embryos (Fig. 3C, lower blot). Taken together with the immunofluorescence data, we conclude that the rate of accumulation of AbdA is specifically reduced in *RpL39^A44^* embryos and we suggest that the resulting lower AbdA levels are insufficient for proper A-P axis patterning and SGP behaviour.

### Reduction in AbdA protein levels are not caused by a reduction in abdA mRNA levels

To test whether the reduction in AbdA levels observed in *RpL39^A44^* embryos results from reduction in *abdA* mRNA levels we first examined the activity of the *abdA* promoter using a transgene in which the *abdA* promoter is fused to the lacZ coding sequence (*abdA>lacZ*; Shimell et al., 1994). By immunofluorescence and quantitative Western blotting, we did not observe a difference in the levels or pattern of expression of LacZ between *RpL39^A44^* and control embryos (Fig. 3D-F). We then examined *abdA* mRNA expression pattern and level directly using *in situ* hybridization and RT-qPCR. The *abdA* expression pattern was identical in wild type and *RpL39^A44^* embryos (Fig. 3G-J). There was a 30% reduction in *abdA* mRNA levels in *RpL39^A44^* embryos (Fig. 3K) as measured by RT-qPCR, however this is not sufficient to fully explain the scale of the reduction in AbdA protein levels.

### N-terminal domain impairs translation of AbdA in RpL39^A44^ mutants

We next wanted to identify whether the critical feature within the *abdA* mRNA responsible for the reduction in AbdA levels in *RpL39^A44^* embryos lies in the coding sequence or the 51 or 31 UTRs. We compared the ability of embryos from wild type and *RpL39^A44^* mothers to express AbdA from an HA-tagged *abdA* construct in which the 51 and 31 UTRs were substituted with those from *hsp70* and *tubulin,* respectively (*UAS abdAHA;* Fig. 4A) using the early zygotic driver (*nulloGal4*). Using an anti-HA antibody, ectopically expressed AbdA was detectable in wild type embryos, however in embryos from *RpL39^A44^* mothers, ectopic AbdA levels were severely reduced (Fig. 4B). As a control, we tested expression of an unrelated HA-tagged gene (*tim17b2*) which was cloned into the same UAS construct and inserted at the same genomic location as the *abdA* construct. We found no difference in the expression level of this protein between embryos from control and *RpL39* mothers (Fig. 4B). We conclude that the critical feature that causes reduced AbdA levels in *RpL39^A44^* embryos lies in the AbdA coding sequence (CDS).

**Figure 4:**
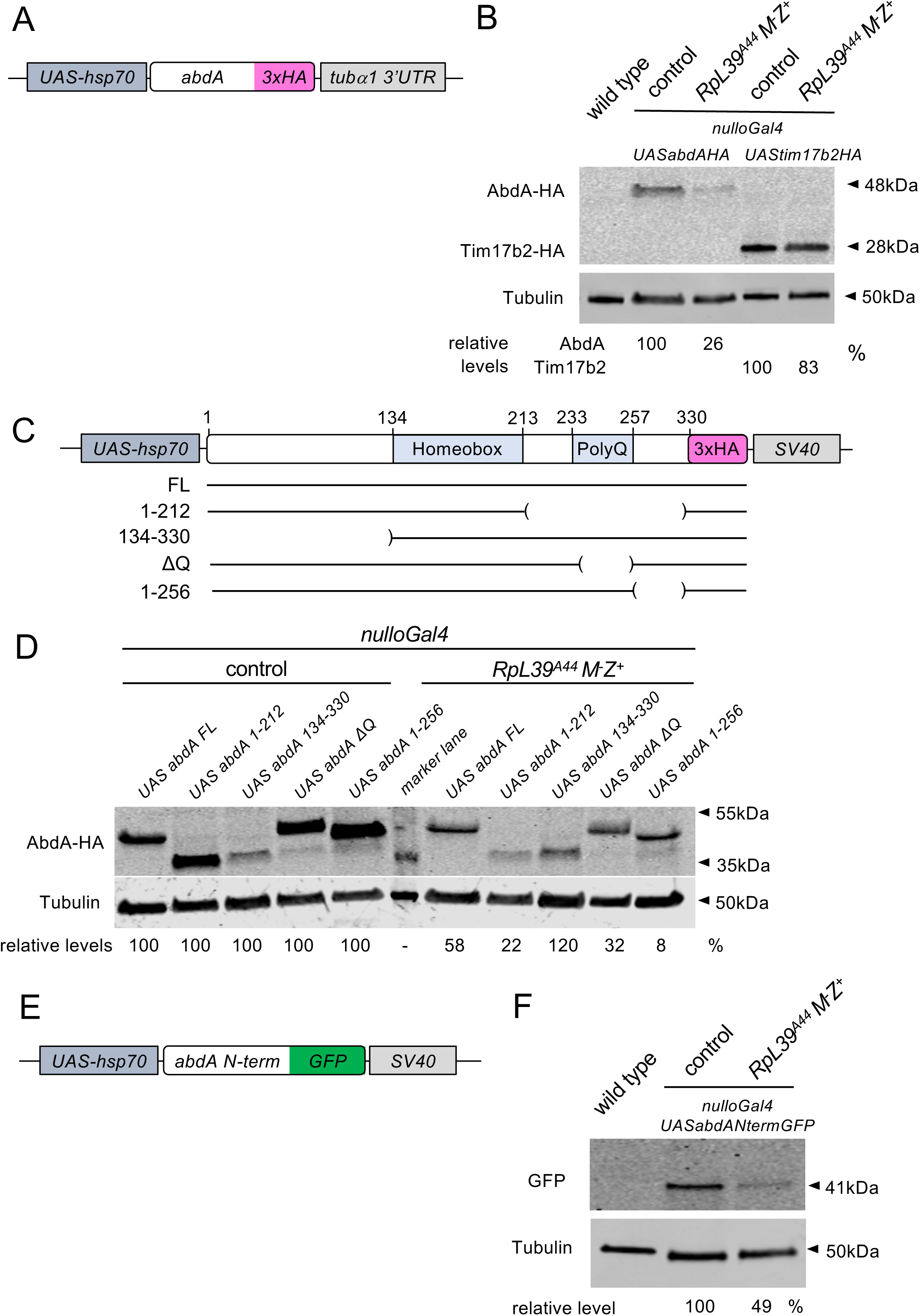
The N-terminus of AbdA is responsible for reduced AbdA levels in RpL39 embryos. (A) Representation of *UAS abdAHA* construct which contains exogenous 5’ and 3’ UTRs. (B) Western blot of 2-4hr embryo extracts from wild type flies or *nulloGal4* (control) and *RpL39^A44^ nulloGal4* mothers mated to *UAS abdAHA* or *UAS tim17b2HA* males and probed with anti-HA and anti-tubulin antibodies. Relative levels indicate AbdA or Tim17b2 band intensity normalized using the tubulin loading control and shown as a percentage relative to the band intensity of the control embryos. Mean levels of AbdAHA and Tim17b2-HA in embryos from *RpL39^A44^* mothers was 27% (n=7, s.d.=13%) and 100% (n=3, s.d.=18%) respectively compared to their expression in the controls. (C) Representation of the deletion constructs showing the portion of the AbdA protein they encode. FL denotes the full length protein and remaining constructs encode proteins with the residues shown. The DQ protein omits 17Q-H-6Q residues. (D) Western blot of 2-4hr embryo extracts from *nulloGal4* and *RpL39^A44^ nulloGal4* mothers mated to *UAS abdAHA* deletion construct males and probed with anti-HA and anti-tubulin antibodies. Relative levels indicate AbdA band intensity normalized using the tubulin loading control and shown as a percentage relative to the band intensity of that construct in the control embryos. Blot shows reduced expression of all constructs except *abdA 134-330* in *RpL39^A44^* embryos. Mean levels of AbdA 134-330 in embryos from *RpL39^A44^* mothers was 89% (n=4, s.d.=36%) compared to in controls. (E) Representation of *UAS abdA N-term GFP* construct. (F) Western blot of 2-4hr embryo extracts from wild type (*w*) files and *nulloGal4* (control) and *RpL39^A44^ nulloGal4* mothers mated to *UAS abdA N-term GFP* males and probed with anti-GFP and anti-tubulin antibodies showing reduced expression of AbdANterm-GFP in *RpL39* embryos. Mean level of AbdAN-termGFP in embryos from *RpL39 nulloGal4* mothers was 41% (n=3, s.d.=14.7%) compared to controls.

The AbdA protein contains a central homeobox domain, and a short poly-glutamine stretch (Q_17_) towards the C-terminus, but is otherwise unannotated (Karch et al., 1990). To narrow down the region(s) responsible for impairing AbdA protein expression, we generated UAS constructs carrying various HA-tagged dissections of the *abdA* CDS (Fig. 4C) and compared the relative protein expression level of each between wild-type and *RpL39^A44^*embryos, again using the *nulloGal4* driver (Fig. 4D). The full-length AbdA construct and three constructs deleting various parts of the C-terminal domain (1-212, ΔQ and 1-256) exhibited reduced expression in embryos from *RpL39^A44^* mothers compared to controls indicating that these retained the feature causing low expression in *RpL39^A44^* embryos. However, while the 134-330 construct lacking the N-terminus exhibited reduced expression in wild-type embryos compared to the other truncations, the levels in both the wild-type and *RpL39^A44^* embryos were similar (Fig. 4D). Together, these results suggest that the compromised expression of AbdA in *RpL39^A44^* is due to a feature of the N-terminal region.

We next tested whether the AbdA N-terminal region is sufficient to reduce expression *RpL39^A44^* background by attaching the coding sequence for AbdA residues 1-133 onto the 51 end of the sequence for EGFP (Fig. 4E). The expression of the resulting N-terminal fusion protein showed reduced expression in embryos from *RpL39^A44^* mothers compared to controls (Fig. 4F). We conclude that the AbdA N-terminal region is sufficient to cause poor translation in a *RpL39^A44^* background.

### Mutation in RpL39 does not affect bulk translation rates

The effects of mutation in *RpL39* could be the result of a general reduction in protein synthesis rates or due to effects that are limited to specific proteins. To distinguish between these possibilities, we looked at the length of the posterior scutellar bristles. A common feature of Minute mutants, caused by mutations in many ribosomal proteins, is a dominant reduction in length of adult bristles caused by an inability to satisfy the heavy protein synthesis demands associated with bristle formation. Whilst we were able to detect significantly shorter posterior scutellar bristles in heterozygous *RpS13* and *RpL14* adult females compared to wild type, as has previously been reported (Marygold et al., 2007), the bristles of both *RpL39^A44^* homozygotes and heterozygotes were not significantly different to wild type (Fig. 5A, B). These data are consistent with mutation in *RpL39* not causing a significant effect on general protein synthesis rates.

**Figure 5:**
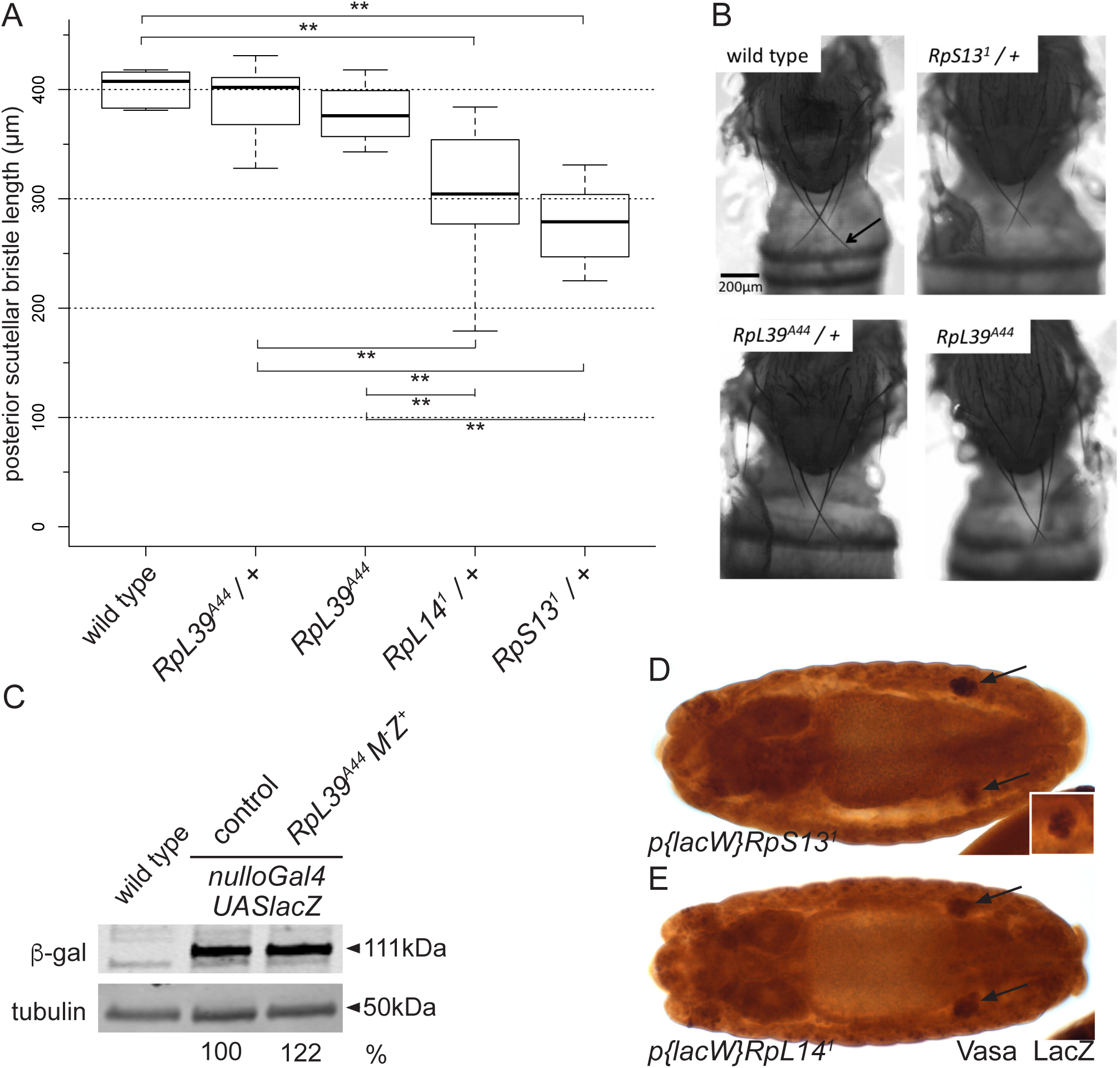
Mutation in RpL39 does not affect bulk translation. (A) Box and whisker plot showing the length of the posterior scutellar bristles of *RpL39^A44^* homozygous and heterozygous adults compared to heterozygous mutants in other ribosomal proteins. There was a statistically significant difference between genotypes as determined by one-way ANOVA (F(4,45) = 21.87, p = < .0001). Significant differences as determined by post hoc comparisons using the Tukey HSD test are indicated (** P ≤ 0.01) whereas other differences were not significant (n=10 bristles from 5 adult females for all genotypes). (B) Thoracic region of *Drosophila* adults showing the pair of posterior scutellar bristles (arrow) for some of the genotypes in (A). (C) Western blot of 2-4hr embryo extracts from wild type flies (*white*) and *nulloGal4* (control) and *RpL39^A44^ nulloGal4* mothers mated to *UAS lacZ* males and probed with anti-β-galactosidase and anti-tubulin antibodies. Relative levels indicate β-galactosidase band intensity normalized using the tubulin loading control and shown as a percentage relative to the band intensity in the control embryos. Blot shows no reduced expression of β-galactosidase in *RpL39^A44^* embryos. Mean levels of β-galactosidase in embryos from *RpL39^A44^* mothers was 230% (n=3, s.d.=153%) compared to in controls. (D-E) Dorsal view of a stage 15 embryo homozygous for *lacZ* containing P-element insertional mutations in *RpS13* (D) and *RpL14* (E) stained with an antibody against Vasa to highlight the germ cells and LacZ to distinguish homozygous and heterozygous embryos. Background staining is due to LacZ expression from P-element insertion. Arrows point to embryonic gonads. Inset shows gonad that is in different focal plane in (D). Images are representative of 11 and 19 embryos observed for each genotype respectively.

To distinguish between the two hypotheses more directly, we examined the ability of *RpL39* mutant embryos to over-express an unrelated protein, namely β-galactosidase, using the *nulloGal4* driver. There was no reduction in the levels of β-galactosidase between control embryos and *RpL39* maternal mutants (Fig. 5C) indicating that the general rate of protein synthesis is not reduced in *RpL39* mutants.

Conversely, if *RpL39* mutants were causing a general reduction the protein synthesis rate we would expect that Minute mutants would phenocopy *RpL39* mutants. Therefore, we examined gonad formation in *RpS13* and *RpL14* mutants. We found that germ cells migrated to the gonad and clustered as in wild type in homozygous *RpS13* and *RpL14* mutant embryos (Fig. 5D, E compared to Fig. 1C). We conclude that Minute mutants do not phenocopy *RpL39* mutants and therefore a general reduction in protein synthesis rates is not sufficient to cause the patterning defects associated with *RpL39*.

### RpL39 interacts with components of the Nascent polypeptide-Associated Complex but not Xrp1

To investigate the mechanism of how mutation in *RpL39* is affecting AbdA levels we performed a candidate-based screen for genetic interactors of *RpL39^A44^* using the same logic and mating schemes as Fig. 2E. Given the location of RpL39 near the peptide exit tunnel of the ribosome, we focused on various ribosome-associated chaperones as well as checkpoint proteins that become activated upon ribosome dysfunction.

The NAC complex binds reversibly to the ribosome and interacts with nascent chains aiding their co-translational folding and preventing premature interaction with the Signal Recognition Particle (SRP) (reviewed in Deuerling et al., 2019). NAC is a heterodimer of α and β subunits and both contact a number of ribosomal proteins near the exit tunnel, including RpL31 (Pech et al., 2010). Given that RpL31 genetically interacts with RpL39 in yeast (Peisker et al., 2008) we tested whether mutations in the *Drosophila* NAC α and β subunits, encoded by the *nacα* and *bicaudal* (*bic*) genes respectively, interact with *RpL39^A44^.* For *nacα* we tested a P element insertion into the *nacα* 51 UTR, which reduces its mRNA levels (Braat et al., 2004). When homozygous *RpL39^A44^*females were mated to males carrying the *nacα* insertion in trans to a dominantly marked balancer chromosome, the percentage of adult offspring that emerged inheriting the mutant chromosome was significantly reduced compared to the controls in which *white* females were used (Fig. 6A), indicating a strong genetic interaction. For *bicaudal* (*bic*) we used the *bic^1^* allele, a spontaneous mutation that lies 31 to the *bic* coding sequence (Markesich et al., 2000). This allele showed no interaction with *RpL39^A44^* (data not shown).

**Figure 6:**
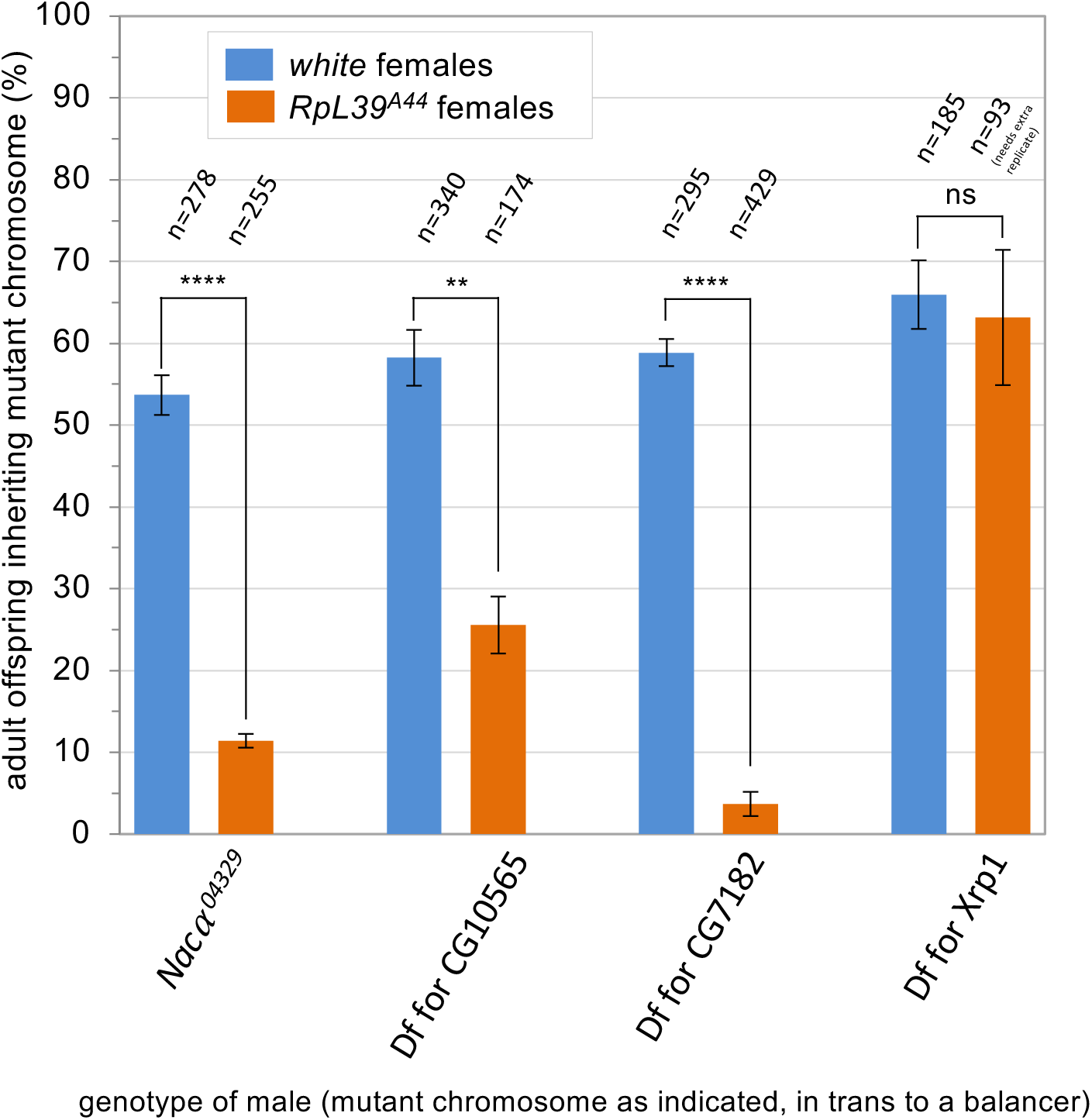
RpL39 interacts with alleles and deficiencies uncovering components of the nascent polypeptide-associated complex (NAC) and ribosome-associated complex (RAC) but not Xrp1. Graph showing the percentage of offspring inheriting the mutant chromosome from type of cross depicted in Fig2E. NAC complex: Naca and *bic*; RAC complex: Df(3L)BSC553 used for DNAJC2 orthologue CG10565; Df(3L)BSC815 used for HSPA14 orthologue CG7182; and Df(3R)Exel6181 used for Xrp1. Significance shown based on Student’s t test (ns P > 0.05; * P ≤ 0.05; **P ≤ 0.01; *** P ≤ 0.001; **** P ≤ 0.0001).

The RAC complex is a conserved heterodimer consisting in *S.cerevisiae* of Zuo1 (in humans DNAJC2) and the ATPase dead Hsp70-type protein Ssz1 (in humans HSPA14) that recruit and stimulate the ATPase activity of the Hsp70-type ATPase active chaperones Ssb1 and Ssb2 to promote co-translational protein folding (Leidig et al., 2013). Zuo1 has only a single *Drosophila* homologue, CG10565. The gene encoding the *Drosophila* orthologue of Ssz1 has not been formally assigned because, as a member of a large Hsp70 family, it has multiple *Drosophila* homologues. Phylogenetic analysis indicates that the *Drosophila* orthologue of Ssz1 is CG7182 (Fig. S4A). Inspection of the positions of ATPase critical sites in the alignment of Ssz1 with CG7182 indicates it, like all Ssz homologues identified to date, is unable to hydrolyse ATP due to the absence of these crucial residues (Fig. S4B, Leidig et al., 2013). This further suggests functional conservation between Ssz1 and CG7182. CG10565 and CG7182 have been poorly studied and no well characterised alleles are available. We therefore tested whether deficiencies that uncover these genes (but which will also include neighbouring genes) interact with *RpL39^A44^*. We found that deficiencies that uncover both CG10565 and CG7182 showed a strong genetic interaction with *RpL39^A44^* (Fig. 6A).

We tested whether a known interactor of other Rp mutants, Xrp1, also interacts with *RpL39^A4^.* Xrp1 is a putative transcription factor that is necessary for the cell competition that eliminates clones of cells mutant for several Rps including *RpS3, RpS12, RpL36* and *RpL19* (Baillon et al., 2018; Ji et al., 2019; C. H. Lee et al., 2018). We found no interaction between *RpL39* with a deficiency that removes the Xrp1 gene indicating that the mechanism by which *RpL39* is affecting AbdA levels is Xrp1 independent (Fig. 6A).

## Discussion

Here we have shown that a P24S mutation of *Drosophila* RpL39 causes an embryonic anterior-posterior patterning defect due to reduced protein levels of one particular hox protein, AbdA. We speculate that the mechanism involves a failure to relay the need for ribosomally-associated chaperone complexes that help certain nascent peptides to fold. This is based on the position of RpL39 adjacent to the peptide exit tunnel of the ribosome (Fig. 7) (Hopes et al., 2022; Nissen et al., 2000; Zhang et al., 2013), and the genetic interactions of RpL39 with deficiencies that uncover ribosome-associated chaperones (Fig. 6). This hypothesised failure to induce their binding may be constitutive or one that would normally be induced based on interactions between RpL39 residues and nascent peptides. Whether this causes translation to stall or the protein to fold incorrectly and be degraded remains an open question.

**Figure 7:**
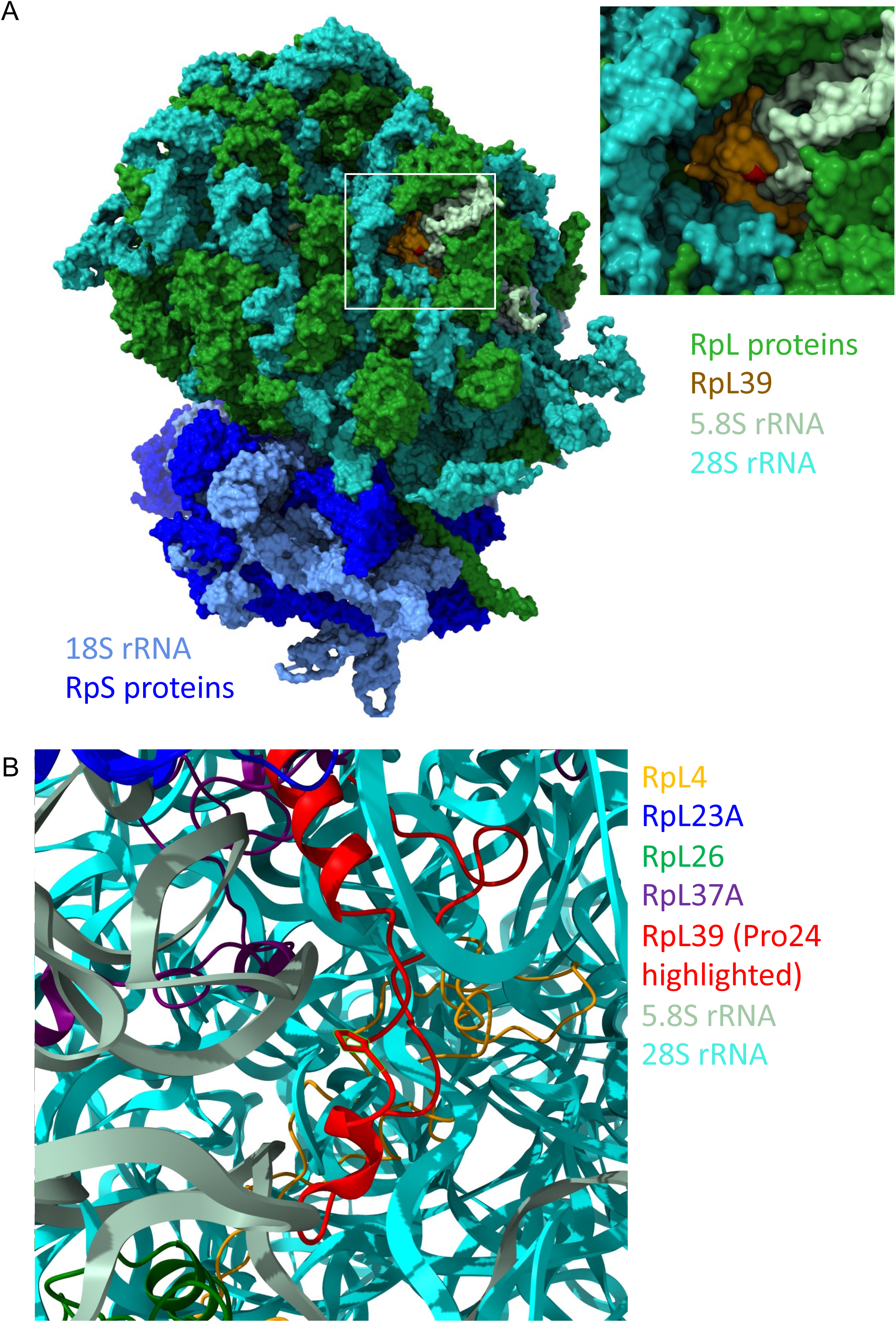
RpL39 location within the Drosophila ovary 80S ribosome. Structure of *Drosophila* 80S ribosome (PDB: 6xu8; (Hopes et al., 2022) showing (A) small subunit (lower part) and location of RpL39 in the large subunit (upper part). Inset shows RpL39 and the proline 24 residue (red) mutated in *RpL39^A44^*. (B) Cartoon diagram of RpL39 and the proline 24 residue in the context of surrounding proteins and rRNA.

These data are consistent with results in mouse. RpL39L knockout embryonic stem cells showed reduced stability of a subset of proteins enriched for long α-helical domains, supporting a role for this tunnel environment in their efficient folding (Banerjee et al., 2024). Knockdown of RpL39 in embryos did not affect overall protein synthesis rates but did activate unfolded protein response pathway genes which is suggestive of protein folding defects (S. J. Lee et al., 2025)

RpL39 is essential in *Drosophila,* as knockdown leads to larval lethality (López-Varea et al., 2021), therefore, the P24S substitution, though in a conserved residue, is not likely to be a null allele as *RpL39^A44^* mutants are homozygous viable. This is consistent with modelling the effect of the substitution using AlphaFold, which did not predict a change to the 3D structure of RpL39 (data not shown). The nearest atoms to the sidechain of the mutated proline are those of the 28S rRNA and are more than 5Å away (Fig. 7b) providing some space for substitution. Interestingly, the NAC β subunit N-terminal tail inserts into the ribosomal exit tunnel and, among other interactions, contacts a nearby residue in RpL39, namely the conserved R28 (Gamerdinger et al., 2019). One possibility is that the P24S mutation interferes with how NAC samples the nascent chain of AbdA.

The connection between a *Drosophila* ribosomal protein and embryonic hox gene expression is conserved in mouse. The mouse large subunit protein RpL38 is required for translation of a subset of hox genes during mouse embryogenesis (Kondrashov et al., 2011). Mechanistically, however, the link is quite different: In mouse, it is proposed that the hox genes in question all have an internal ribosome entry site (IRES) that requires RpL38 for efficient translation (Xue et al., 2015). How RpL38 links structurally to the use of IRESs remains an unanswered question and subsequent work has suggested that altered global translation can confer gene-specific effects through altered start codon selection stringency (Ivanov et al., 2022).

A number of conceptually similar situations have been discovered recently. Haploinsufficiency of mouse *Eif3c,* a component of initiation factor complex eIF3 causes Sonic hedgehog (Shh) gain of function phenotypes caused by reduced translation of the Shh receptor and negative regulator of the pathway, Ptch1 mRNA (Fujii et al., 2021). The mechanism here is different again and involves Eif3c binding to pyrimidine rich motifs within the 51 UTR of target mRNAs. Insufficiency of 40S ribosomal proteins RpS26 and RpS25 reduces production of the toxic polyglycine protein FMRpolyG in fragile X–associated conditions, possibly because its CGG-repeat–containing, 51 UTR is especially dependent on these proteins for translation (Tutak et al., 2025).

Gene duplications have led to ribosomal protein paralogues in many species. Whilst *Drosophila* contains a single RpL39 gene, mammalian genomes generally encode a RpL39 paralogue. In mice testis, RpL39 is replaced by its paralogue RpL39L. This testis specific ribosome co-translationally folds a subset of proteins that are involved in sperm formation (H. Li et al., 2022). The authors speculate that the substitution of RpL39 by RpL39L, which differs by only three amino acid substitutions, modifies its charge and hydrophobicity properties, changing the surface properties at the exit tunnel, therefore affecting the co-translational folding of nascent peptides. This is consistent with our hypothesis that specific residues of *Drosophila* RpL39 play a pivotal role in the translation and folding of certain proteins.

One open question is why the *RpL39^A44^* mutation causes a very specific patterning defect, due to reduction in a single hox gene. This specificity is surprising given that in the examples of ribosomal mutants and tissue specific paralogue expression mentioned above, several to many hundreds of proteins are affected (Kondrashov et al., 2011; Li et al., 2022).

We speculate that our RpL39 mutant may affect many more proteins than AbdA. However, the very rapid nature of embryogenesis and limited time window for patterning genes to be expressed, due to the regulatory cascades needed for their regional specific expression, means that some genes may be highly susceptible to lower translation or folding rates during such periods.

Although we found that AbdA mRNA levels were somewhat reduced in *RpL39* embryos (Fig. 3K), this does not seem sufficient to completely account for the decrease in detected protein levels. One possibility is that this mRNA reduction is a symptom of the reduced AbdA expression, rather than a cause. If the *RpL39* mutation leads to prolonged stalling of the ribosome during translation, this could trigger mRNA surveillance pathways such as No-Go Decay (NGD), leading to the premature termination of translation, and degradation of the nascent peptide and mRNA (reviewed in (Monaghan et al., 2023).

We have mapped the region in AbdA that causes the defect in AbdA levels to the N-terminal region (residues 1-133) of the protein. This region contains motifs (residues 109-118) that allow binding of AbdA with co-factors such as Extradenticle that promote DNA binding specificity (Lelli et al., 2011). The solved *Drosophila* AbdA 3D structure is restricted to the homeodomain (PDB Entry 8F37, residues 137-197), and the AlphaFold full length structure predicts this region to be mostly disordered but with low confidence (Varadi et al., 2024). Further dissection of the N-terminal region will help identify the causative feature or property.

A number of ribosomal proteins are linked to disease such as ribosomopathies, developmental disorders caused by abnormal ribosome synthesis or ribosome dysfunction, as well as contributing to cancer progression. For example, knockdown of RpL39 inhibits the growth of human pancreatic cancer cells (C. odong Li et al., 2014). *RPL39* and its gain-of-function mutation A14V have oncogenic activity in breast cancer (Dave et al., 2014, 2017). Mutations in ribosomal proteins may cause preferential translation of certain mRNAs, creating a pro-oncogenic proteome (Elhamamsy et al., 2022). This work provides a new entry point to understand the mechanisms that underlie these changes.

## Materials and Methods

### Drosophila genetics

The *Drosophila* stocks used in this study are given in table S1. Flies were reared on standard Bloomington recipe food at 25°C. For the genetic interaction studies, crosses were set-up in triplicate or quadruplicate. For the control crosses using *white* females, a single vial was scored. For *RpL39^A44^* females, due to the low number of offspring they produce, the vial was flipped twice and the offspring in all three vials was scored.

### Embryo staining and cuticle preparations

Embryos were collected on apple juice agar plates at 25°C. Embryos were dechorionated in 2% sodium hypochlorite (Sigma) and then fixed for 20 mins in 1:4 ratio of 4% formaldehyde in PBS and heptane. The embryos were devitellinized by shaking in 1:1 methanol:heptane and stored in methanol at −20°C. For immunohistochemistry, embryos were hydrated in PBS 0.1% Tween-20 (PBST) and then blocked in 0.5% BSA in PBST. The following antibodies were used: guinea pig anti-AbdA (1:1000; Brian Gebelein); mouse anti-AbdB (1:10; DSHB 1A2E9); mouse anti-Fasciclin III (1:50; DSHB 7G10), Rabbit anti-Vasa (1:10000; Ruth Lehmann); rabbit anti-LacZ (1:10000; Cappel); rat anti-HA (1:500 Roche). Embryos were mounted in Aquamount (Polysciences) and visualized using a Zeiss LSM 880.

Fluorescent and colourimetric *in situ* hybridizations were carried according to (Tripathy et al., 2014) and (Lehmann & Tautz, 1994) respectively. A template for the *abdA* RNA probe was prepared by amplification of embryonic cDNA using abdAprobeF and abdAprobeR primers (table S2) and JumpStart REDTaq (Sigma). The 1.7kb product was TA cloned into the vector pCR®2.1 (Invitrogen). The vector was linearized with KpnI and a DIG labeled RNA probe transcribed using T7 RNA polymerase.

Cuticle preparations were carried out according to (Alexandre, 2008).

### Vector construction

pUASt AbdA-HA vectors were constructed via a pUASt-3xHA-attB vector. The latter was generated via PCR amplification of a 3 x haemagglutinin (HA) tag from genomic DNA prepared from *M{UAS-abd-A.ORF.3xHA.GW}ZH-86Fb* (flyORF – F000044) flies using the primers in table S2, and cloned into pUASt-attB (Bischof et al., 2007) linearised with XhoI and XbaI using the In-Fusion HD cloning kit (Takara Bio). The various AbdA fragments were PCR amplified from genomic DNA prepared from *M{UAS-abd-A.ORF.3xHA.GW}ZH-86Fb* flies using the primers in table S2 and cloned into the pUASt-3xHA-attB vector linearised with XhoI and NotI.

For the pUASt AbdA N-term GFP construct, the *abdA* N terminal region (corresponding to residues 1-133) and GFP were PCR amplified from genomic DNA prepared from *M{UAS-abd-A.ORF.3xHA.GW}ZH-86Fb* and *UAS GFP S65T* flies respectively. The resulting fragments were cloned into pUASt-attB linearised with XhoI and XbaI using the In-Fusion HD cloning kit.

The resulting constructs were integrated into chromosome II via microinjection into line *vas-int; attP40* (University of Cambridge Fly Facility).

### Western blotting

Embryos were collected on apple juice agar plates at 25°C. Where necessary, the embryos were aged at 25°C before harvesting. Embryos were dechorionated as above and stored at - 80°C. Embryos were homogenized in 30μl of lysis buffer (20mM Tris, 1mM EGTA, 320mM sucrose, 0.1% Triton X-100, 1mM NaF, 10mM β-glycerophosphate, 1% Sigma protease inhibitor cocktail) per mg of embryos, solubilized using 6x solubilization buffer (24% SDS; 30% glyercol; 30% β-mercaptoethanol; 2.5% bromophenol blue; 1.5M TrisHCl), boiled at 95°C for 5 minutes and insoluble material was pelleted by centrifugation. The soluble fraction was separated on a TGX 4-20% gradient gel (Roche) and blotted to Protran nitrocellulose membrane (Whatman). The following antibodies were used: guinea pig anti-AbdA (1:3000; Brian Gebelein); mouse anti-Tubulin (1:5000; Millipore DM1A); rabbit anti-LacZ (1:20000; Cappel); mouse anti-LacZ (1:2000; Promega); mouse anti-Actin (1:5000; Sigma A1978); mouse anti-Antp (1:60; DSHB 8C11); rat anti-HA (1:2000 Roche); chicken anti-GFP (1:10000 Abcam); 800CW and 680RD IRDye secondaries (1:10000; LI-COR Biosciences). Bands were detected using an Odyssey fluorescence imaging system (LI-COR Biosciences) and band intensities quantitated and normalised using Image Studio (LI-COR Biosciences).

### Reverse transcription-qPCR assays

Embryos were collected and dechorionated as described above, and snap frozen at −80°C. Total RNA was prepared using Trizol/Chloroform, and purified using a RNeasy Mini Kit (QIAGEN) (adapted from Ponton et al, 2011). Any contaminating gDNA was digested with EzDNase (Thermo Fisher), and reverse transcription reactions were carried out in 20 µl total volume using Superscript IV VILO Master Mix (Thermo Fisher). RT-qPCR reactions [4.5 µl cDNA sample (diluted 1:100), 100 nM of each primer, 5 µl PowerUp SYBR Green Master Mix (Thermo Fisher)] were plated in triplicate on 96-well plates (StarLab) and ran on a 7500 Fast RT-PCR System (Applied Biosystems) using the default Fast thermocycling conditions with ROX as a passive reference dye. The qPCR primers used are listed in the Supplementary Material. No-template and no-reverse transcriptase controls were ran alongside the experimental samples. Relative levels of AbdA expression were determined using the comparative CT method, normalised to the relative expression of actin.

## Acknowledgements

This work was funded by a Wellcome Trust Seed Award in Science (205951/Z/17/Z). We thank Ruth Lehmann for hosting the *Drosophila* germ cell migration screen that generated the A44 mutant and Hiroko Sano and Prabhat Kunwar for being part of the screening team. Author contributions: MM, RT, JB, MW and ADR performed the experiments and analysed the data. MM and ADR wrote the manuscript.

