## Supplemental Methods for "A ribosome exit tunnel component is required for robust expression of the *Drosophila* hox gene, *abdominal A*"

**Supplemental Materials and Methods**

**Cloning of *A44***

The *A44* mutant line was isolated from the ethyl methanesulphonate (EMS) mutagenesis screen described in (Barbosa et al., 2007). The gonad formation/germ cell migration phenotype is maternal and is present in all of the offspring laid by females containing germline clones made using the FRT element at cytological location 42B on the right arm of chromosome 2. This places the causative mutation to the right of position 42B. The original line was homozygous lethal but through recombination mapping with a *FRT c px sp* chromosome the lethality and causative mutation were separated, generating viable mutant chromosomes. In this first round of recombination mapping, the causative mutation mapped to the right of the visible marker *plexus* at position 58E4 and to the right of *l(2)tid* at 59F6 by SNP analysis (Fig. S1). In a second round of SNP mapping using recombination with an *If* chromosome the mutation mapped to the left of the gene *nervy* at position 60C5 (Fig. S1). The region between these two positions is 582kb and contains 130 predicted genes. Complementation testing with deficiencies was used to narrow down this region further. The mutation was not uncovered by the deficiencies *Df(2R)BSC136*, *Df(2R)BSC356* or *Df(2R)BSC780*. These deficiencies uncover the majority of this region but fail to uncover a small region at 60B7 which contains the genes *eIF-5A, RpL12, RpL39, Rap21* and *yki.* Whole genome sequencing done in parallel to these experiments (described in Tripathy et al., 2014) identified a single SNP that would cause a coding sequence change in only 1 of these 5 genes, namely a G to A change at position 2R:24,062,235 (release 6) causing a Pro24Ser change in RpL39.

**Phylogenetic analysis of RAC orthologues in *Drosophila***

Homologues of the yeast and human RAC components were identified in *Drosophila* through BLAST searches. A phylogenetic tree was inferred using the NGPhylogeny.fr suite {Lemoine:2019io} to create a multiple sequence alignment using MAFFT (version 7.407, PAM120 substitution matrix) which was trimmed using Gblocks (version 0.91.1) to retain 373 (35%) of the original 1052 positions and used for tree inference by Maximum likelihood using PhyML (version 3.1) with 100 bootstraps.

Barbosa, V., Kimm, N., & Lehmann, R. (2007). A Maternal Screen for Genes Regulating Drosophila Oocyte Polarity Uncovers New Steps in Meiotic Progression. Genetics, 176(4), 1967–1977. https://doi.org/10.1534/genetics.106.069575

Tripathy, R., Kunwar, P. S., Sano, H., & Renault, A. D. (2014). Transcriptional regulation of *Drosophila* gonad formation. Developmental Biology, 392(2), 193–208. https://doi.org/10.1016/j.ydbio.2014.05.026
