## Supplemental figures for "A ribosome exit tunnel component is required for robust expression of the *Drosophila* hox gene, *abdominal A*"

**Figure S1: Recombination and deficiency mapping of A44.**

(A) Depiction of deficiencies uncovering regions between cytological location 59D to 60F on chromosome 2R. Deficiencies listed all complemented *A44* by phenotype thus *A44* mutation is not present within these deficiencies. Recombination mapping places *A44* between the genes *l(2)tid* and *nvy*.
(B) Depiction of genes at cytological location 60B7 between *l(2)tid* and *nvy* that are not uncovered by deficiencies.
(C) Coding sequence change identified by next generation sequencing in RpL39.

**Figure S2: *RpL39<sup>A44</sup>* genetically interacts with *abdA* in gonad formation.**

- (A-B) Maximum intensity Z projections of Vasa stained embryos from *RpL39<sup>A44</sup>* mothers mated to *abdA<sup>46.51</sup>* in trans to a *lacZ* expressing balancer chromosome. Embryos inheriting the *abdA<sup>46.51</sup>* chromosome (A) show no obvious germ cell clusters at the normal gonad position whereas embryos inheriting the balancer chromosome (B) show germ cell mis-migration but gonad clusters are often apparent (arrows). (C) Quantitation of number of germ cell gonad clusters per embryo in embryos from the above cross. Distribution is significantly different (Fisher's exact test $p < 0.005$).

**Figure S3: AbdB expression is expanded in *abdA* mutant embryos but AbdA expression is unaffected in *ubx* mutant embryos**

- (A-D) Lateral views of stage 11 sibling control (A, C) and *abdA<sup>46.51</sup>* (B) *ubx<sup>1</sup>* (D) embryos stained with antibodies against AbdA (A-D) and AbdB (A', B'). Whilst AbdB is mis-expressed in the AbdA domain in an *abdA* mutant embryo (B') there is no expansion of AbdA in *ubx* embryos (D).

**Figure S4: CG7182 is the *Drosophila* homologue of the yeast and human Hsp70-like RAC component.**

(A) Multiple sequence alignment of the *S. cerevisiae* RAC component Ssz1 (NP\_011931.2) and human RAC component HSPA14 (NP\_057383.2) with canonical Hsp70 proteins from *S. cerevisiae* Ssb1 (NP\_010052.1), human HSPA1A (NP\_005336.3) and Hsp70 proteins from *Drosophila* CG7182 (NP\_001286979.1), CG2918 (NP\_001162645.1), Hsc70-1 (NP\_524063.1), Hsc70-2 (NP\_524339.1), Hsc70-3

(NP\_001285139.1), Hsc70-4 (NP\_001262586.1), Hsc70-5 (NP\_001286414.1), HSP70Ab (NP\_524798.2), Hsc70Cb (NP\_001189103.1), Hsp68 (NP\_524474.1). The catalytic centre of Hsp70 proteins contains three elements (E1, E2 and E3) crucial for ATP hydrolysis. The RAC components Ssz1 and HSPA14 binds but are unable to hydrolyse ATP due to mutations in several catalytically important residues (highlighted in red for canonical Hsp70s). *Drosophila* CG7182, CG2918 and HSc70Cb also lack a complete set of these catalytic residues and are predicted to be unable to hydrolyse ATP.

(B) Consensus phylogenetic tree of sequences listed above compiled using maximum likelihood with bootstrap support (%) based on 1000 trees. Identified RAC components in *S. cerevisiae* and humans, Ssz1 and HSPA14 respectively, are highlighted.

Figure S1

A

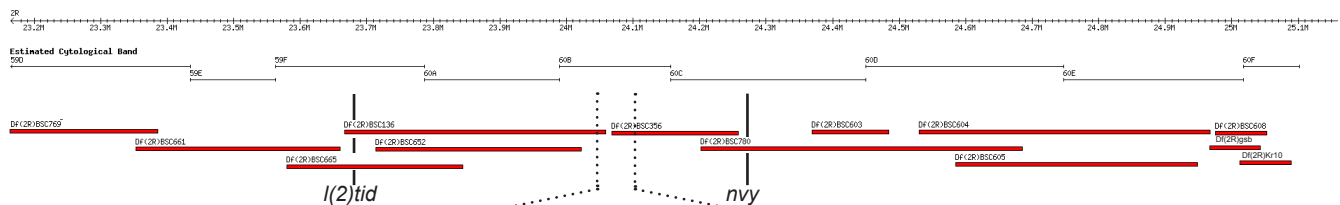

B

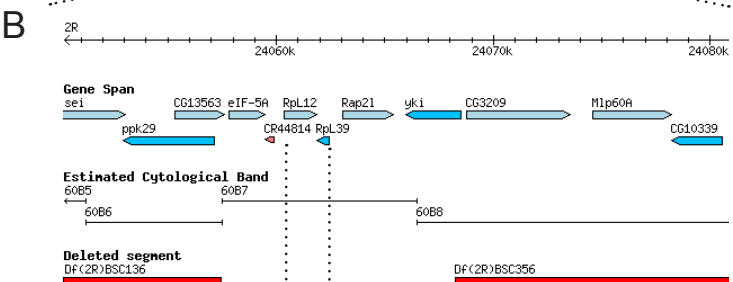

C

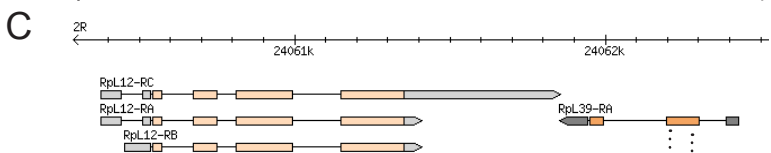

D

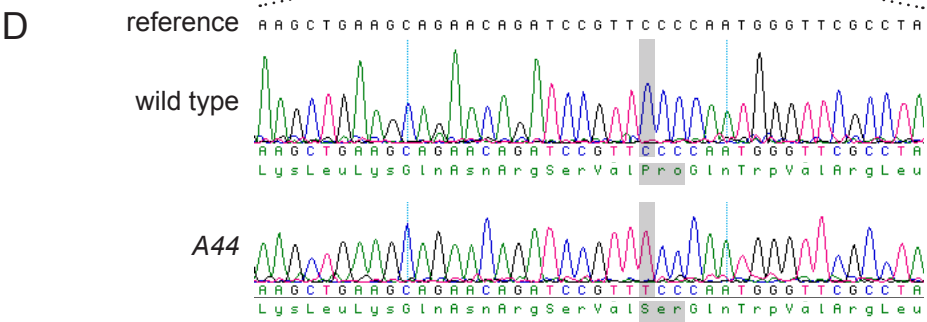

Figure S2

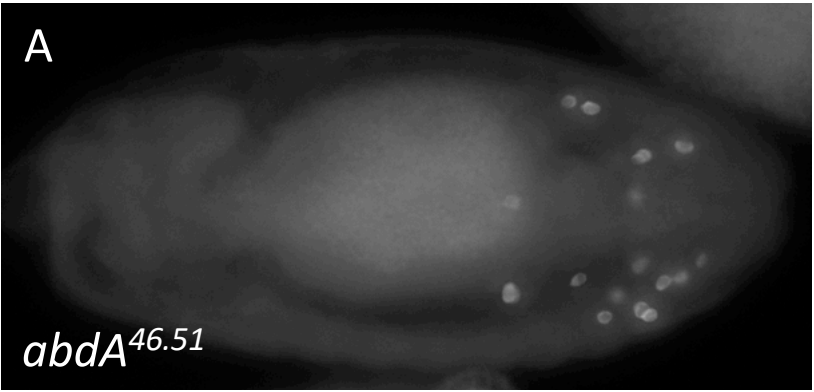

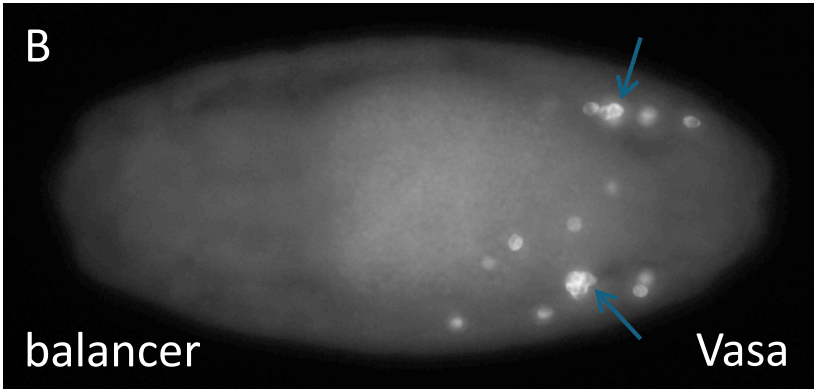

| C | Number of germ cell gonad clusters |  |  |
| --- | --- | --- | --- |
|  | 0 | 1 | 2 |
| *abdA*<sup>46.51</sup> | 15 | 3 | 0 |
| balancer | 2 | 7 | 14 |

Figure S3

*abdA*<sup>46.51</sup> / +

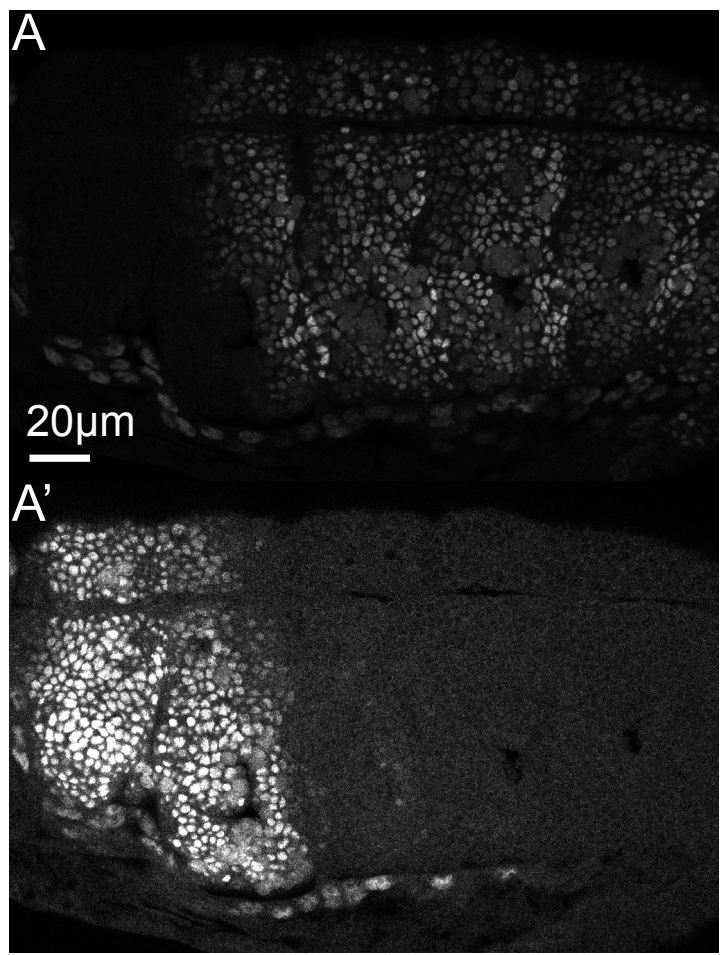

*abdA*<sup>46.51</sup>

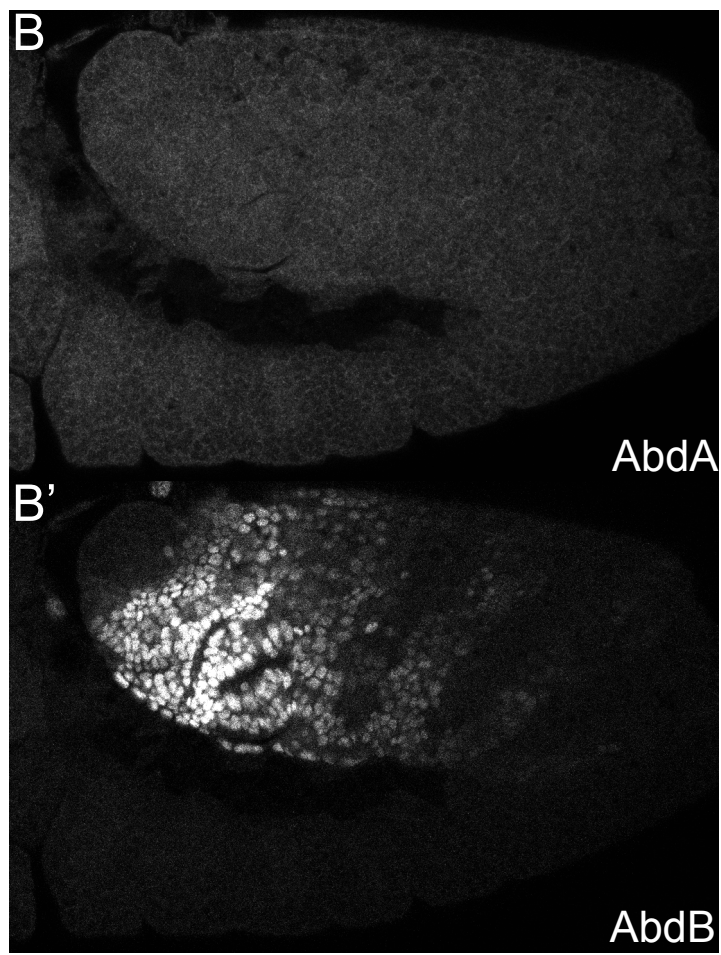

*ubx*<sup>1</sup> / +

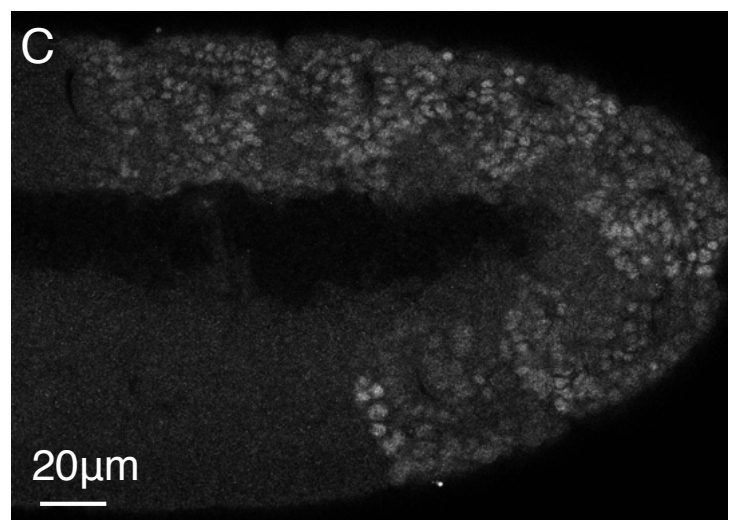

*ubx*<sup>1</sup>

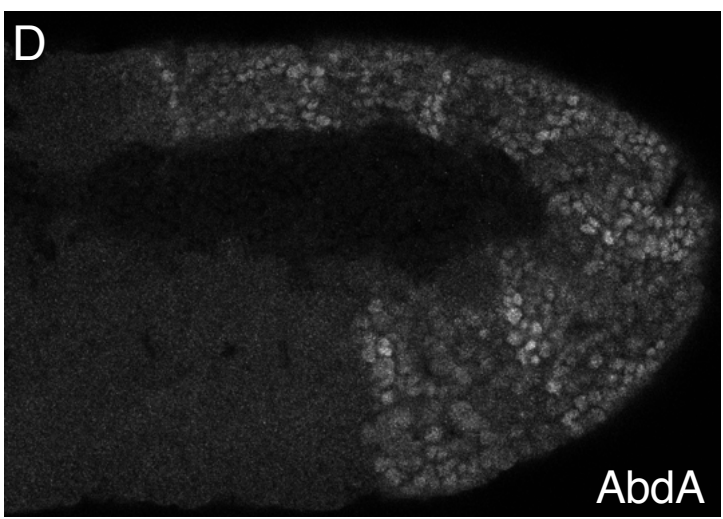

Figure S4

A

|  |  | E1 |  | E2 |  | E3 |  |
| --- | --- | --- | --- | --- | --- | --- | --- |
| Sc_SSZ1 | 8 | ITFGNTSS | 15 | 70 | FRDF | 73 | 200 GGIRSD 205 |
| Hs_HSPA14 | 6 | VHLGCTSA | 13 | 67 | VKQI | 70 | 199 GGTSLS 204 |
| Dm_CG7182 | 7 | IKIGNSTL | 14 | 70 | SFQL | 73 | 215 GGLYSD 220 |
| Dm_CG2918 | 25 | VDLGSEWM | 32 | 87 | LLDL | 90 | 221 GAYKTS 226 |
| Sc_SSB1 | 12 | IDLGTTYS | 19 | 72 | AKRL | 75 | 204 GGGTFD 209 |
| Hs_HSPA1A | 9 | IDLGTTYS | 16 | 70 | AKRL | 73 | 201 GGGTFD 206 |
| Dm_Hsc70-1 | 9 | IDLGTTYS | 16 | 70 | AKRL | 73 | 201 GGGTFD 206 |
| Dm_Hsc70-2 | 9 | IDLGTTYS | 16 | 70 | AKRL | 73 | 201 GGGTFD 206 |
| Dm_Hsc70-3 | 34 | IDLGTTYS | 41 | 96 | AKRL | 99 | 226 GGGTFD 231 |
| Dm_Hsc70-4 | 9 | IDLGTTYS | 16 | 70 | AKRL | 73 | 201 GGGTFD 206 |
| Dm_Hsc70-5 | 57 | IDLGTTNS | 64 | 119 | TKRL | 122 | 246 GGGTFD 251 |
| Dm_HSP70Ab | 6 | IDLGTTYS | 13 | 67 | AKRL | 70 | 198 GGGTFD 203 |
| Dm_Hsc70Cb | 6 | IDFGNESC | 13 | 67 | FKRL | 70 | 202 GHSSLQ 207 |
| Dm_Hsp68 | 6 | IDLGTTYS | 13 | 67 | AKRL | 70 | 198 GGGTFD 203 |

B

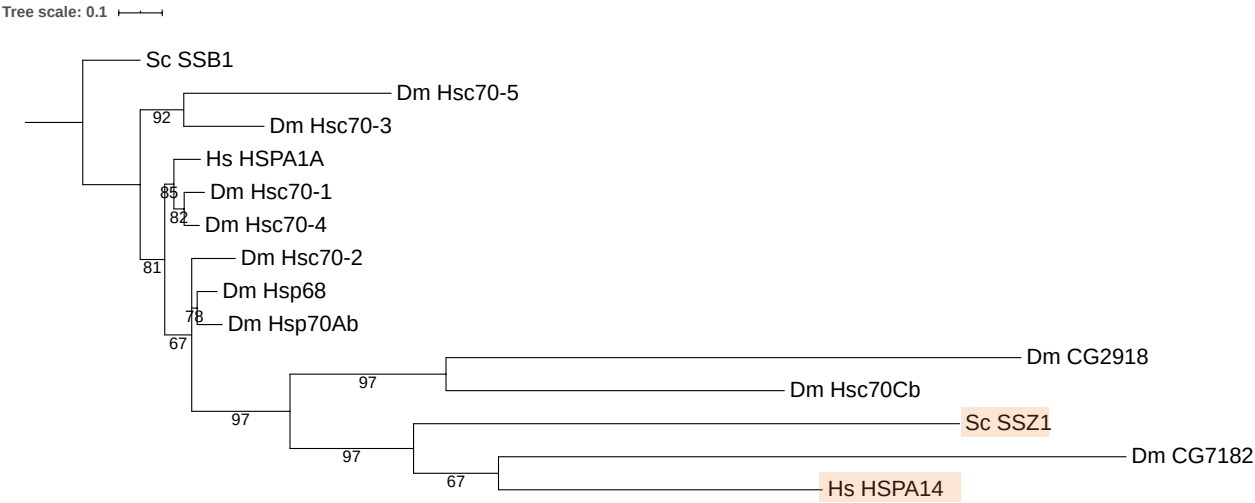
